# Identification of ICAM-1–targeting DNA aptamers as a host-directed strategy to inhibit Human Rhinovirus infection

**DOI:** 10.64898/2026.04.20.717810

**Authors:** Jessica Dellavedova, Chiara Campera, Silvia Ancona, Monica Rebecchi, Vittorio Panzeri, Thomas Carzaniga, Luca Casiraghi, Stefano Rocca, Stefano Di Ciolo, Alessandro Pedretti, Claudio Tirelli, Marco Buscaglia, Tommaso Bellini, Alessandra Romanelli, Alessandro Villa, Electra Brunialti, Elisa Borghi, Paolo Ciana

## Abstract

Exacerbations of respiratory viral infections significantly contribute to morbidity and healthcare burden. Among these viruses, Human Rhinoviruses (HRVs) are the most frequent causative agents of upper respiratory tract infections. To date, over 150 HRV serotypes have been identified, classified into three species: HRV-A, HRV-B, and HRV-C. No antiviral therapies are currently available against this viral family, largely due to the high serotype diversity and limited cross-protection. The major group of HRVs relies on the Intercellular Adhesion Molecule-1 (ICAM-1) receptor to infect airway epithelial cells, making ICAM-1 an attractive target for broad-spectrum therapeutic interventions. Here, we report the development of nucleic acid-based aptamers designed to disrupt ICAM-1–HRV binding and thereby prevent viral infection. Aptamers are single-stranded DNA molecules that fold into precise three-dimensional structures, enabling highly specific protein recognition. Using a Systematic Evolution of Ligands by EXponential Enrichment (SELEX) approach guided by a minimal peptide mimicking the ICAM-1 viral binding interface, a library of >10^24^ random single-stranded DNA sequences was screened. Through iterative rounds of selection, we identified eight candidate 77-nt DNA aptamers, which were subsequently evaluated for their potential using *in silico* and *in vitro* assays, as well as functional assays in human epithelial cells. From this strategy, two lead aptamers were selected that effectively inhibited HRV-A16 replication in a concentration-dependent manner, as measured by viral titers (TCID₅₀ assay) and viral RNA quantification by RT-PCR. These findings demonstrate the potential of ICAM-1-targeting aptamers as antiviral agents capable of preventing HRV entry. By targeting a host receptor and creating a protective barrier at the cell surface, this approach may offer a broadly applicable strategy against multiple HRV serotypes, paving the way for the development of novel antiviral interventions.

**Graphical abstract:** 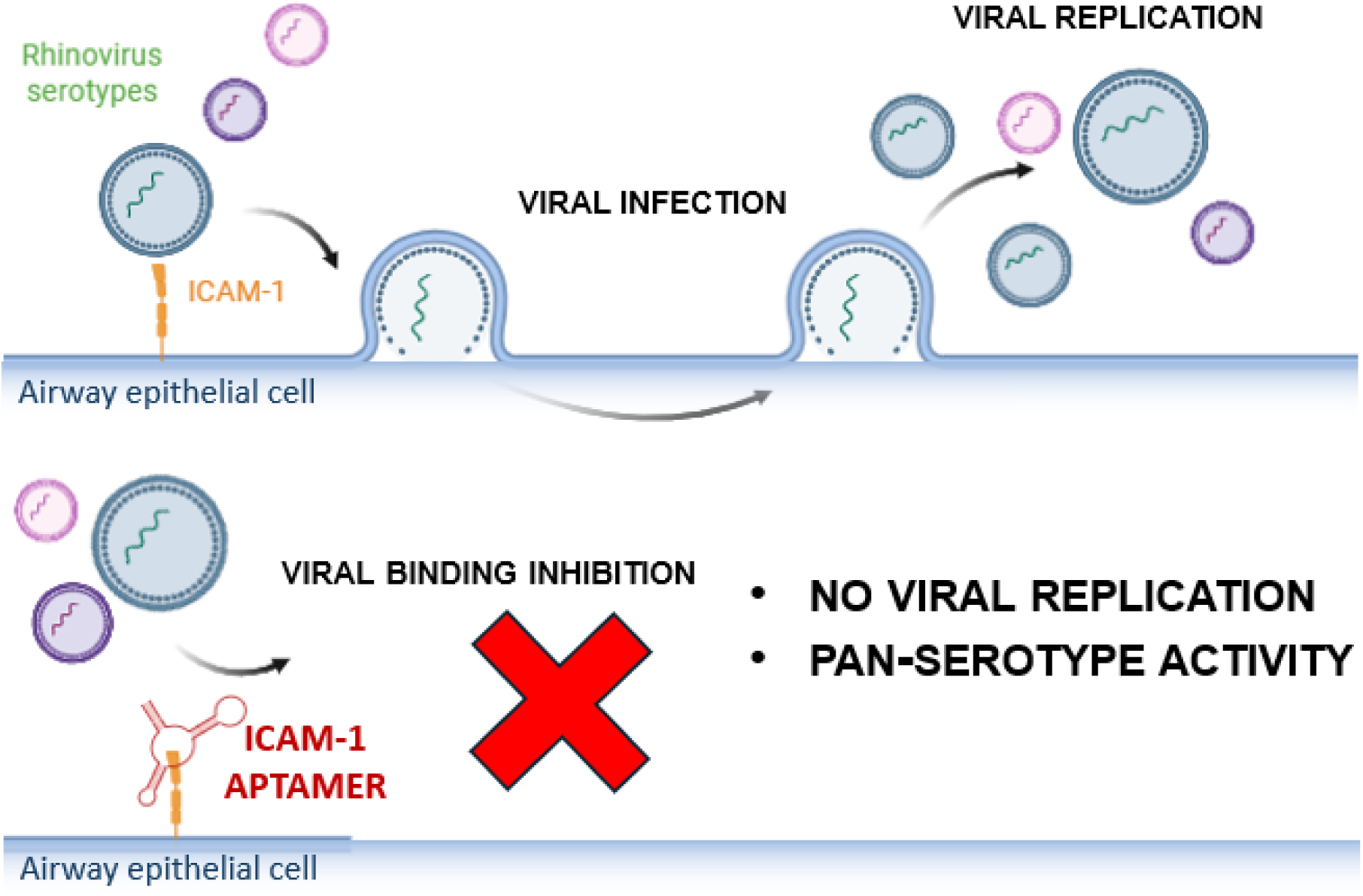

## Introduction

Rhinoviruses (HRVs) are among the most common respiratory viruses in humans and represent the primary cause of upper respiratory tract infections such as the common cold [1,2]. HRV infections are highly prevalent across all age groups, with children experiencing an average of 6-10 infections per year and acting as a major reservoir for viral transmission within the community [3–5]. Although they are typically self-limiting and resolve without medical intervention, HRVs can cause severe disease in vulnerable populations, including infants and individuals with underlying chronic respiratory conditions such as asthma or chronic obstructive pulmonary disease (COPD) [4,6,7]. In these patients, rhinoviral infection is a frequent driver of acute exacerbations often leading to complications such as secondary pneumonia and hospitalization [8,9]. Notably, HRVs have been detected in up to 60% of asthma exacerbations [10], underscoring their critical role in disease morbidity. In addition to their clinical impact, HRVs contribute substantially to healthcare utilization and socioeconomic burden worldwide [11]. Despite this impact, no approved antiviral treatments or vaccines are currently available. The development of effective therapeutics has been consistently hampered by the extensive genetic diversity of HRVs, which comprise over 150 serotypes, classified into three species (A, B and C) [12]. This diversity results in limited cross-protective immunity and renders traditional vaccine strategies largely ineffective [13]. An alternative promising strategy to circumvent viral variability is targeting host factors essential for viral entry. HRVs enter host cells by engaging specific receptors: while minor group viruses utilize cadherin-related family member 3 (CDHR3) or member of the low-density lipoprotein receptor (LDLR) family, the majority of serotypes belonging to species A and B (the “major group”) utilize Intercellular Adhesion Molecule-1 (ICAM-1) [14,15]. While this approach has been validated by the clinical success of agents like the CCR5 antagonist maraviroc [16] and the anti-CD4 monoclonal antibody ibalizumab [17], previous attempts to target ICAM-1 using monoclonal antibodies or soluble receptor decoys have faced significant hurdles, including high production costs, potential immunogenicity, and limited stability in the respiratory environment [13,18]. To overcome these limitations, nucleic acid aptamers have emerged as a promising alternative [19]. Building on our previous work where we successfully developed a DNA aptamer targeting the ACE2 receptor to block SARS-CoV-2 infection [20], we hypothesized that a similar host-directed approach could be applied to the HRV-ICAM-1 interaction. Aptamers are short, single-stranded DNA or RNA oligonucleotides that fold into defined three-dimensional structures, enabling them to bind targets with high affinity and specificity [21,22]. Identified through the *in vitro* process known as SELEX (Systematic Evolution of Ligands by EXponential enrichment), this strategy allows precise control over selection conditions and enables the enrichment of ligands targeting specific functional epitopes, representing a key advantage over conventional approaches. In this study, we employed SELEX to identify DNA aptamers specifically targeting the distal extracellular domains of ICAM-1 critical for viral attachment. By shifting the focus to the conserved host receptors rather than the rapidly mutating viral capsid, this strategy aims to achieve broad-spectrum inhibition of major-group of HRVs. Here, we characterized the binding affinities and structural features of these aptamers and demonstrated their efficacy in inhibiting HRV infection and reducing the production of infectious virus in a cell-based model. This approach highlights the potential of host-directed aptamers as a versatile platform for the development of next-generation antiviral therapeutics.

## Results

### Identification of a minimal ICAM-1 peptide for the SELEX procedure

To identify a peptide suitable as a bait for SELEX procedure and capable of mimicking the viral interaction region, a structure-guided analysis of HRV-ICAM-1 complex was performed using the available PDB crystal structure [23]. Residue-level interaction energy decomposition revealed specific segments contributing most significantly to complex stabilization. In agreement with previous studies [23,24], the analysis highlighted the HRV binding interface surrounding the critical residue Lysine 29 (K29) (Fig. 1A). To enable the selection of aptamers targeting this interface, short peptide fragments were designed to mimic the ICAM-1 regions directly involved in HRV interaction. Two initial candidates (peptide 1 and 2) were identified based on favourable interaction energies (Fig. 1B). However, peptide 2 was excluded due to low binding efficiency and unfavourable net charge (-1), while peptide 1, although energetically favourable, was considered too long for SELEX applications. Therefore, shorter overlapping segments (∼20 amino acids) derived from peptide 1 were further evaluated. This approach led to the identification of eight additional peptides (Fig. 1B). Candidate selection was guided by the following criteria: (i) total interaction energy, (ii) binding efficiency (energy per residue), (iii) number of positively charged residues, and (iv) inclusion of the key residue K29 within the sequence. Although peptide 3 displayed the highest binding efficiency, peptide 4 (from now named ICAM-1 29–50) was prioritized due to its enriched content of positively charged residues, anticipated to improve electrostatic complementarity with nucleic acids. To assess whether ICAM-1 29-50 preserves features of full-length ICAM-1, molecular dynamics simulations were performed (Fig. S1 and Table S1). The peptide displayed a dynamic conformational profile in solution, indicative of structural flexibility, yet it recurrently sampled conformations resembling the crystallographic state, indicating that native-like structures remain accessible. Overall, ICAM-1 29-50 retains key structural and physiochemical features of the ICAM-1 binding interface, supporting its use as a conformational mimic of the HRV-ICAM-1 binding interface and was therefore chosen as bait for the SELEX process.

**Fig. 1.**
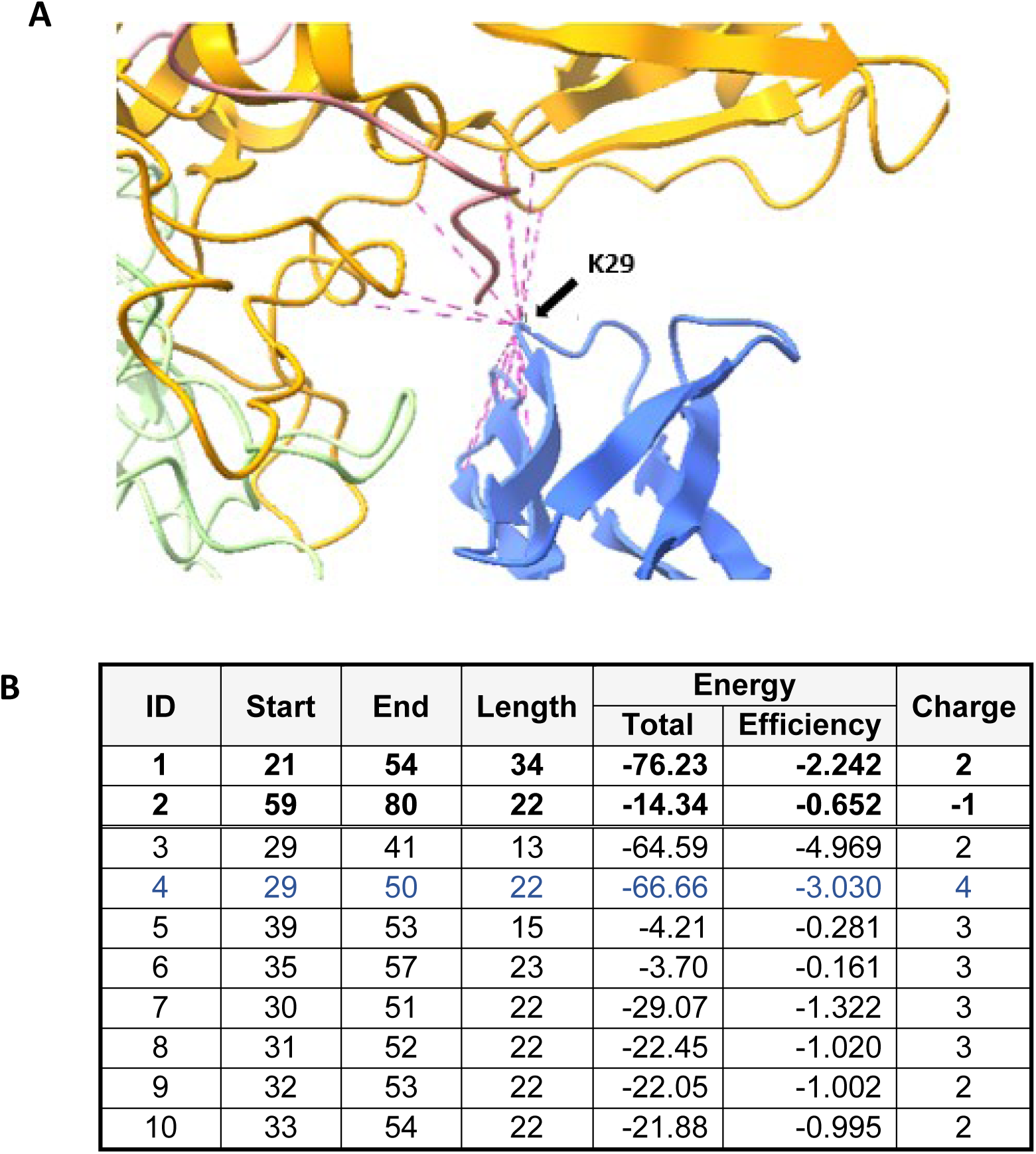
Structure-based selection of ICAM-1-derived peptides targeting the HRV binding interface. A. Representative model of rhinovirus capsid protein VP1 (yellow) and VP2 (green) in complex with human ICAM-1 (blue), obtained from Protein Data Bank database (1D3E). Interacting residues at the interface are shown in pink, highlighting ICAM-1 K29 as a key residue mediating the interaction. B. Peptides identified from PDB:1D3E following energy minimization and interaction energy decomposition at the residue level. For each peptide, the start and end positions within the ICAM-1 sequence, peptide length, total interaction energy (Total, kcal/mol), binding efficiency (Efficiency, kcal/mol/residue), and net charge are reported.

### Aptamer selection through SELEX procedure

Aptamers recognizing the ICAM-1 29-50 peptide were isolated through an iterative SELEX procedure following standard protocols [25,26]. Briefly, the peptide was synthesized and purified by high-performance liquid chromatography–mass spectrometry (HPLC-MS) and functionalized with a N-terminal biotin to enable immobilization on streptavidin-coated beads, which served as the binding substrate during selection. A library of >10²⁴ random single-stranded DNA sequences was used as the starting material. The selection consisted of an initial round of positive selection against ICAM-1 29-50, followed by a counter-selection (negative selection) step using uncoated beads to eliminate non-specific binders, and subsequently nine additional rounds of positive selection. A total of 10 positive selection cycles were performed (Fig. 2A), and amplified pools were subjected to next-generation sequencing at rounds 4, 6, 8, 9 and 10, leading to the identification of eight enriched sequence named I1-I8 (Fig. 2A). To evaluate their binding specificity, each sequence was chemically synthesized and tested for binding to ICAM-1 29–50-coated beads, using uncoated beads (empty) as a negative control (Fig. 2B). Sequences I5 and I8 showed a statistically significant preference for peptide-coated beads compared to empty beads, followed by I4 and I7, whereas the other sequences did not exhibit specific ICAM-1 29-50 binding.

**Fig 2.**
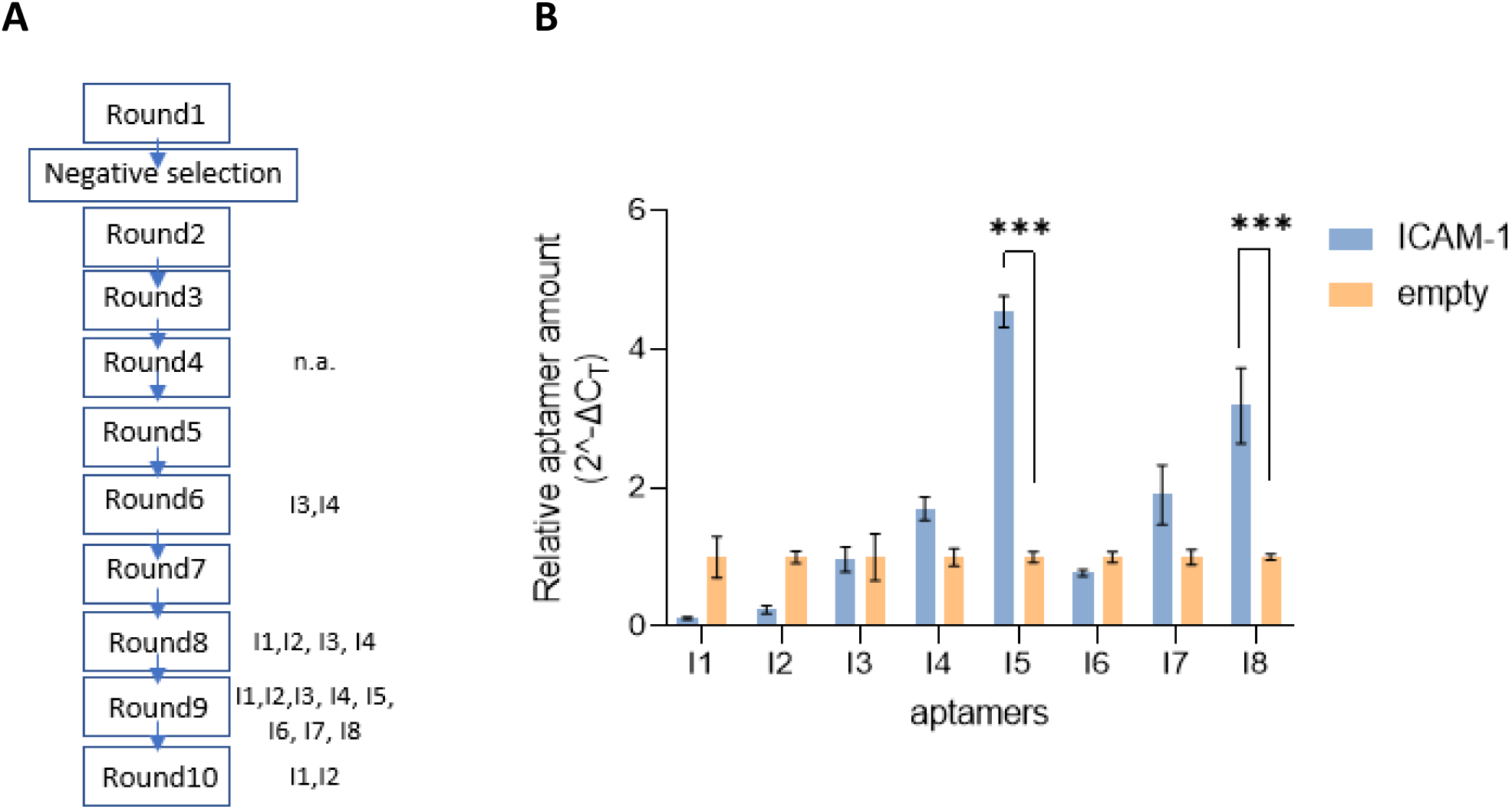
Identification of aptamers with high affinity for ICAM-1 peptide. A. Scheme of aptamer selection using the ICAM-1 29–50 peptide. Enriched sequences identified by next-generation sequencing at each round are indicated (I1-I8). No enriched sequences were detected at round 4 (n.a., not applicable). B. Evaluation of the specific ICAM-1 29-50 peptide binding ability for each aptamer. Aptamer recovery after the binding assay was quantified by RT-PCR. Bars indicate relative aptamer levels eluted from beads without the peptide (empty) or peptide-coated beads (ICAM-1) calculated using the 2^-ΔCT^ method. Mean ± SEM, n = 3 replicates. Statistical analysis was performed using 2way ANOVA, followed by Sidak’s multiple comparisons test vs empty beads.

### In silico and in vitro validation

To further validate the binding properties of the identified aptamers, both *in silico* and *in vitro* analyses were performed. First, to gain structural insight, three-dimensional models of the aptamers were generated and subjected to molecular docking [25] against the full-length ICAM-1 protein. The resulting complexes displayed variability in predicted binding energies, buried surface areas (BSA), and energetic contributions including desolvation energy and internal energy, indicating distinct binding modes among the different sequences (Table 1). Interaction efficiency, defined as binding energy normalized by BSA, ranged from −0.0978 to −0.1740 (a.u./Å²), with I1 showing the highest value (−0.1740), followed by I4 (−0.1532), I8 (−0.1458), and I5 (−0.1316). A combined scoring function integrating binding energy, internal energies, desolvation energy and BSA was then applied to rank the complexes. Based on this final score, sequences I1, I5, I8 and I4 emerged as the most favourable binders, with scores of -0.075, -0.072, -0.065 and -0.062, respectively. I1 showed the highest interaction efficiency, consistent with a highly optimized local interface, although associated with relatively small BSA. In contrast, I5 and I8 displayed more balanced energetic profiles, combining favourable binding energies with low desolvation penalties and moderate energy differences between free and bound states. I4 exhibited the most favourable binding energy (-276 a.u.), although associated with a larger difference between internal energy terms, suggesting a higher energetic cost upon binding. The remaining sequences (I2, I3, I6 and I7) showed weaker binding energies and less favourable overall scoring, resulting in lower ranking.

**Table 1.**
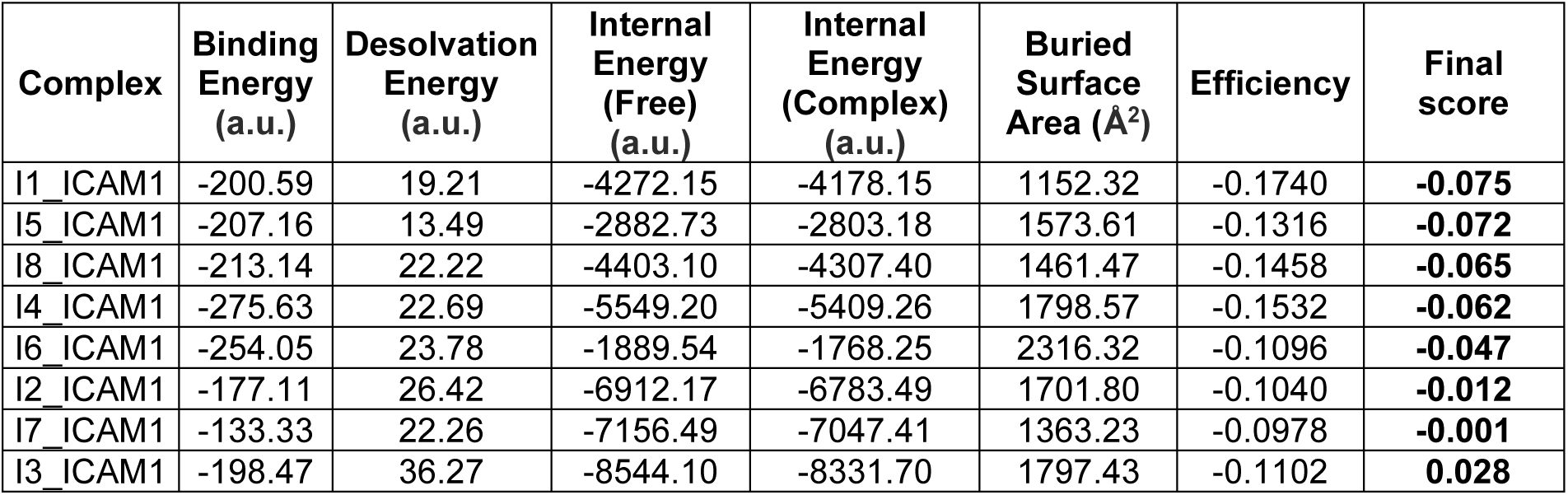
HADDOCK docking analysis of aptamer–ICAM-1 complexes. Predicted interaction parameters for aptamer–ICAM-1 complexes generated by HADDOCK 2.4 are reported, including binding energy, desolvation energy, internal energy (free and complex states), and buried surface area (BSA). Energies are expressed in arbitrary units (a.u.). Efficiency is calculated as binding energy normalized to interface size, while the final score integrates energetic contributions relative to BSA. Complexes are presented in order from the best to the worst final score.

Next, surface plasmon resonance (SPR) spectroscopy [26] was employed to quantify aptamer–peptide interactions for the most promising candidates (I1, I4, I5, I8). Increasing concentrations of chemically synthesized aptamers were injected over a Biacore sensor chip functionalized with the ICAM-1 29-50 peptide, and dissociation constants (K_D_) were determined (Fig. 3). Consistent with preliminary binding screening (Fig. 2B), sequence I1 exhibited weak binding, as indicated by markedly higher K_D_ value, confirming its limited interaction with ICAM-1 peptide. In contrast, I5 and I8 displayed significantly lower K_D_ values, indicative of high-affinity binding and supporting their selection as lead candidates, while I4 showed intermediate affinity.

**Fig. 3.**
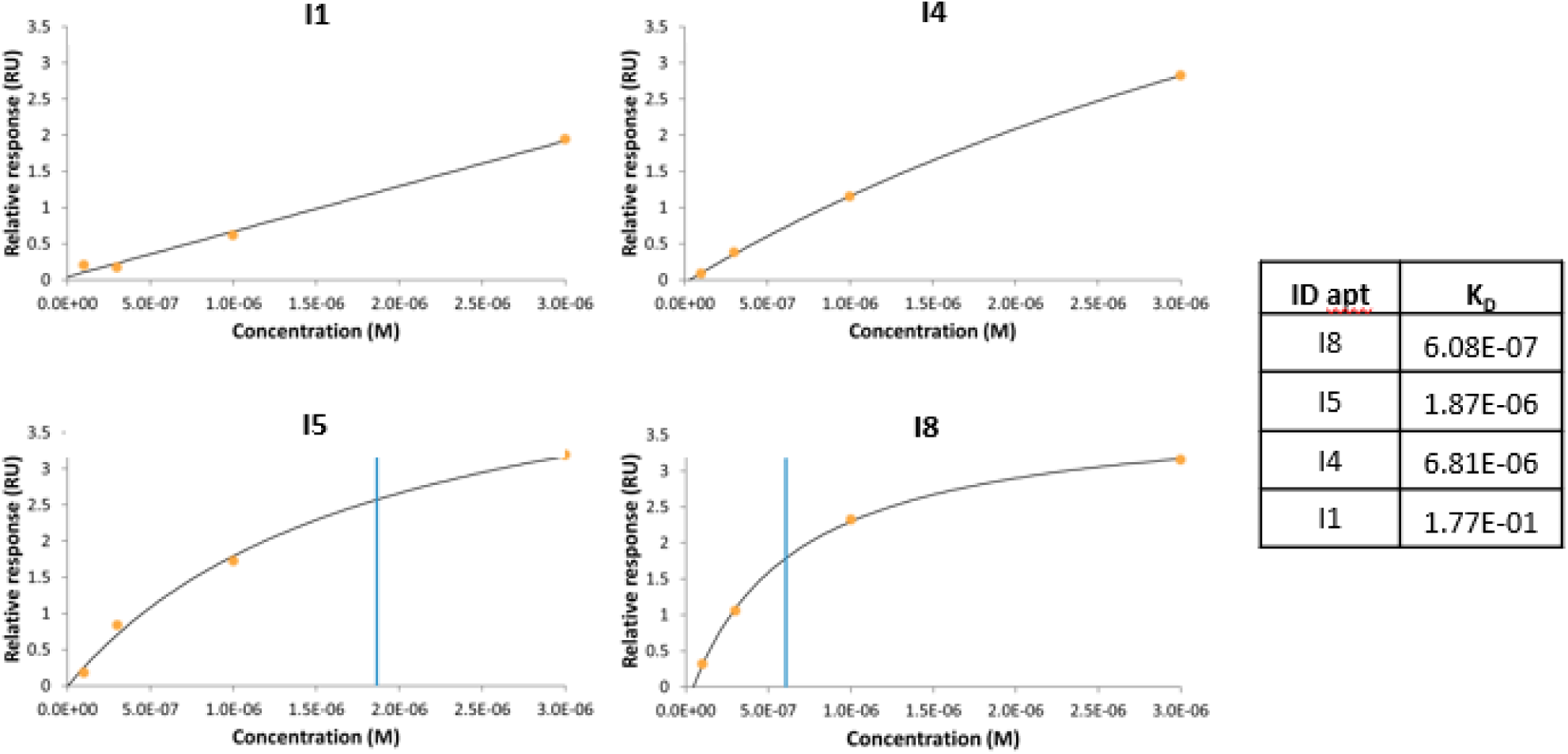
Surface plasmon resonance (SPR) binding curves of aptamer sequences on ICAM-1 peptide. The plot shows binding curves measure by Biacore 1K (Cytiva) for aptamers I1, I4, I5 and I8 as indicated in the panel titles, against the ICAM-1 29-50 peptide. The biotinylated peptide was immobilized on a streptavidin-coated SPR channel. Aptamers were injected at increasing concentrations at a flow rate of 10 µL/min in PBS buffer at 25°C. Binding responses at equilibrium (orange dots) were fitted with sigmoidal curves (black lines). The apparent dissociation constants (K_D_) are indicated by vertical blue lines and summarized in the table.

### ICAM-1 targeting aptamers inhibit HRV-A16 infection

Having established I4, I5 and I8 as the strongest ICAM-1 binders, we next investigated whether these aptamers could effectively inhibit HRV infection. To this end, H1-HeLa cells were infected with a commercial HRV-A16 strain at a MOI of 0.1 and after 72 hours incubation antiviral efficacy was determined by measuring both viral titers (TCID_50_ assay) and viral RNA levels (HRV-specific qPCR). (Fig. 4A). To validate the experimental setup, the capsid-binding inhibitor pleconaril was used as a positive control; as expected, it yielded a dose-dependent reduction in viral replication, confirming the robustness of the infection model (Fig. S2) and the dose of 1 µM used as a standard. The antiviral potential of ICAM-1 targeting aptamers was assessed by pre-incubating cells with two different concentration (1 and 0.1 µM) prior to HRV-A16 infection. As shown in Fig. 4, I5 and I8 significantly inhibited HRV-A16 replication at the concentrations tested. In particular, I5 reduced infectious viral titers to ∼10–20% of control levels at 1 μM, with a consistent decrease in viral RNA (∼30%), indicating concordant inhibition across both readouts. Notably, a measurable antiviral effect of I5 was already observed at 0.1 μM, with a partial reduction in both infectious titers and viral RNA levels, suggesting dose-dependent activity. Similarly, I8 induced a strong reduction in both parameters at 0.1 and 1 μM, supporting a comparable inhibitory profile. In contrast, I4 exhibited negligible antiviral activity, with values comparable to vehicle-treated controls. Overall, these findings demonstrate that ICAM-1-targeting aptamers can effectively block HRV-A16 infection, supporting a host-directed antiviral mechanism.

**Fig. 4.**
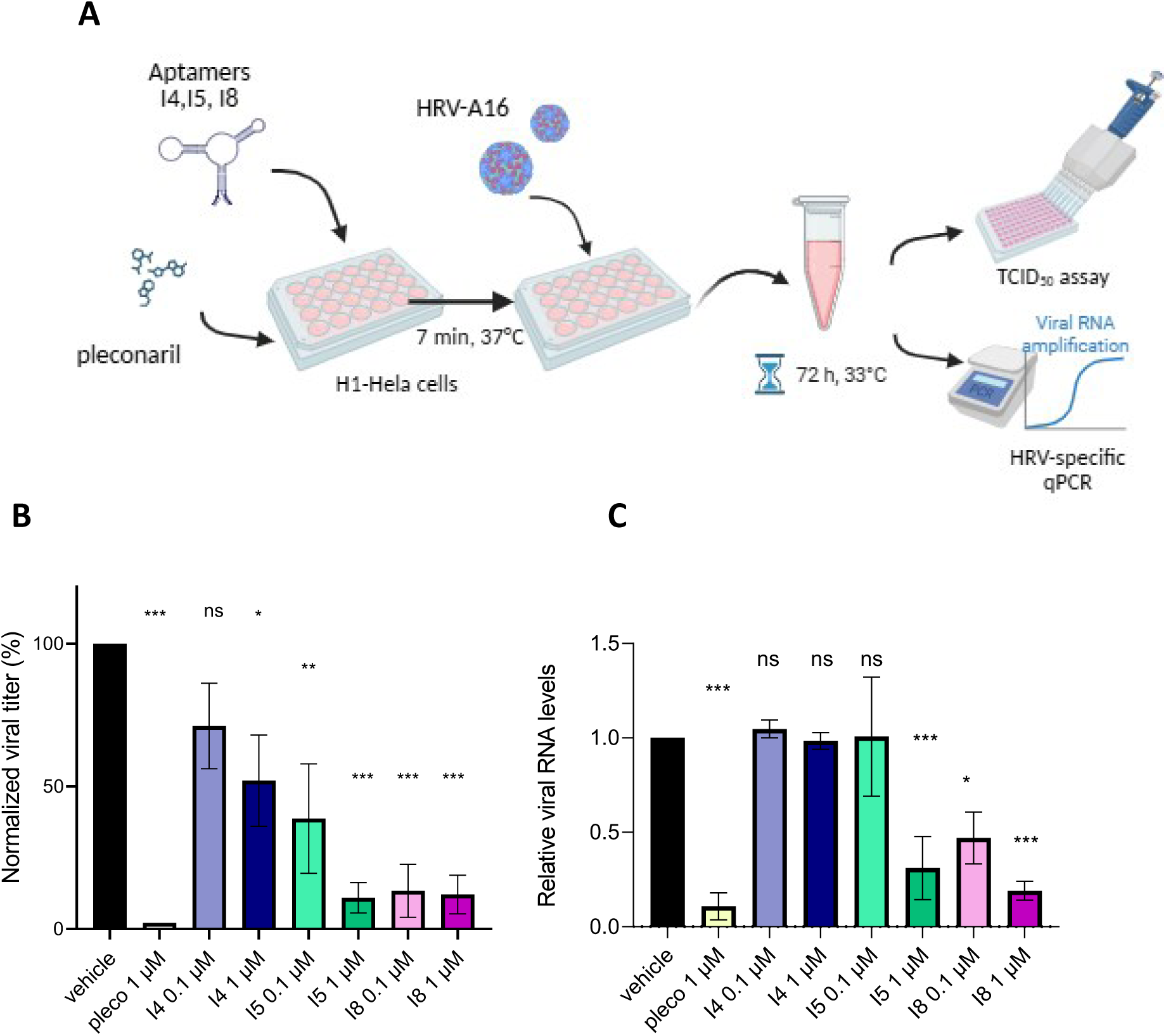
Inhibition of HRV-A16 infection in H1-HeLa cells by selected aptamers. A. Schematic representation of the infection experiment to assess the ability of aptamers to block rhinovirus infection. Aptamers (I4, I5 and I8) were tested at doses of 1 and 0.1 µM. As positive control, for assessing viral inhibition, a VP1 inhibitor (Pleconaril, pleco) was also included. H1-HeLa cells were pre-treated with aptamers for 7 minutes and next the infection with HRV-A16 (MOI 0.1) was performed. After 1 hours, the inoculum was removed and infection was allowed to proceed for 72 hours at 33°C. Infectious virus released into the supernatant was quantified by TCID₅₀ assay (B), and viral RNA levels were measured (C). B. Normalized viral titers (%), expressed relative to vehicle (100%) and mock (0%) (mean ± SEM). C. Viral RNA levels assessed by RT-PCR using HRV-specific primers. Data are shown as 2^-ΔCt^. Statistical significance was determined by ordinary one-way ANOVA followed by Bonferroni’s multiple comparison correction vs vehicle.

## Discussion

In post-pandemic era, it has become increasingly evident that influenza-like respiratory illnesses are caused by a heterogeneous group of viruses belonging to distinct families, each with unique replication strategies and antiviral susceptibilities [27–29]. Despite this complexity, currently available antiviral treatments remain largely virus-specific, limiting their effectiveness and offering little in terms of broad-spectrum protection and pandemic preparedness. Although HRVs have been long considered agents of mild disease, their involvement in severe clinical outcomes, including exacerbations of asthma and COPD [8,30] underscore the need for innovative therapeutic strategies. In this context, we explored a host-directed approach based on DNA aptamers targeting ICAM-1, the primary entry receptor for the major group of rhinoviruses. By focusing on a conserved host factor rather than rapidly mutating viral proteins, this strategy is expected to provide a higher genetic barrier to resistance and potentially broader activity across multiple serotypes. Previous attempts to inhibit ICAM-1-mediated entry have been previously attempted with monoclonal antibodies or decoy receptors [31,32]. However, aptamers offer superior advantages, including high target specificity, lower production costs, and reduced immunogenicity [21,33]. In present work, applying a SELEX procedure we identified potential ICAM-1 binders that were characterized using a combination of computational and experimental approaches. While *in silico* analyses highlighted multiple candidate binders, integration of energetic parameters and interface properties proved essential to discriminate between compact, high-efficiency interactions and more biologically relevant binding modes. In particular, sequences I5 and I8 emerged as the most promising candidates, displaying balanced energetic profiles and consistent binding *in vitro*. Notably, the discrepancy observed for I1, which exhibited high interaction efficiency in docking but weak experimental binding, further emphasizes the limitations of structure-based scoring alone and the importance of experimental validation. Functionally, I5 and I8 significantly inhibited HRV-A16 infection, supporting the feasibility of blocking viral entry through ICAM-1 targeting DNA aptamers. However, several challenges must be addressed to translate these results into clinical applications. A primary limitation of the current study is the use of unmodified DNA aptamers in a simplified cell-based model. Translation to *in vivo* application will require chemical modifications to enhance nuclease resistance and stability [34,35]. Furthermore, given the physiological role of ICAM-1 in leukocyte adhesion and immune cell trafficking, careful evaluation of potential off-target effects and dosing strategies will be necessary to avoid interference with normal immune functions [31]. In conclusion, this work, alongside our previous success in targeting the ACE2 receptor for SARS-CoV-2 [20], reinforces the validity of host-receptor targeting as a versatile antiviral platform through DNA aptamers. Future efforts will focus on evaluating these optimized aptamers against a wider panel of HRV serotypes and in a more physiologically relevant models, such as three-dimensional human airway epithelia. More broadly, this approach may be extended to other viral pathogens that rely on conserved host entry factors, contributing to the development of next-generation, resistance-resilient antiviral therapeutics.

## Materials and methods

### Structure-guided peptide design and molecular dynamics simulations

The crystal structure of the HRV-ICAM-1 complex (PDB 1D3E) was used as the starting model for structural analysis. Missing hydrogen atoms were added using VEGA ZZ [36]. The structure was subsequently optimized using NAMD3 [37] with the CHARMM36m force field [38]. Interaction energies between HRV and ICAM-1 were calculated using non-bonded terms of the force field and decomposed at the residue level to identify key binding regions. Based on this analysis, peptide candidates were selected according to interaction energy, binding efficiency, charge and inclusion of residue K29. Molecular dynamics simulations of the best candidate were performed using NAMD3. The peptide was solvated in an explicit water box, energy-minimized and subjected to equilibration and production runs under standard conditions. Trajectory analysis included calculation of Cα-RMSD, RMSF, and solvent-accessible polar surface area (PSA) to evaluate conformational stability and physicochemical properties. Detailed description of simulation parameters is provided in the Supplementary Materials.

### Synthesis of ICAM-1 peptide and chemical characterization

The ICAM-1 29-50 peptide and its biotinylated analogue were synthesized by microwave-assisted solid-phase peptide synthesis (SPSS) using a Liberty Blue CEM peptide synthesizer. The biotinylated derivative was obtained through on-resin functionalization of the fully assembled peptide. Briefly, an amino hexanoic acid linker was coupled to the N-terminus, followed by conjugation of biotin. This modification was introduced to enable immobilization and facilitate its use in the SELEX procedure. Both peptides were purified by semi-preparative RP-HPLC. Analytical RP-HPLC and mass spectrometry were employed to assess purity and confirm molecular identity. Purified peptides were subjected to multiple lyophilization cycles to remove residual counterions potentially introduced during purification.

### Systematic Evolution of Ligands by Exponential Enrichment (SELEX) procedure

The SELEX procedure was performed using XELEX DNA Core Kit (EURx, Poland, cat. n°: E3650) according to the manufacturer’s instructions and as previously described [20]. The selected ICAM-1 peptide (ICAM-1 29-50) was synthesized with an N-terminal biotin tag and dissolved in nuclease-free water at a final concentration of 3 mg/mL. A total of 80 µg of peptide was immobilized on 2 mg of pre-washed Dynabeads™ MyOne™ Streptavidin C1 (Invitrogen, USA, cat. n°: 65001) by incubating on a rotating shaker for 30 min at room temperature, followed by four washes with PBS pH 7.4. Unoccupied streptavidin binding sites were subsequently blocked with 1 µM free biotin for 5 min. After washing with PBS pH 7.4, ICAM-1-loaded beads were resuspended in SELEX buffer (NaCl 140 mM, KCl 2 mM, MgCl_2_ 5 mM, CaCl_2_ 2 mM, Tris pH 7.4, Tween 20 0.05%) and used for incubation with the ssDNA random library, provided in the kit. The ssDNA library consisted of a 40-nucleotide randomized central region, flanked by two constant 18-bp sequences, required for primer annealing during PCR amplification, as follows: 5’-TGACACCGTACCTGCTCT-40nt randomized sequence -AAGCACGCCAGGGACTAT-3’, corresponding to a theoretical diversity of approximately 10^24^ unique sequences. For the first round of selection, 100 μg of the ssDNA library were resuspended in 500 μL SELEX buffer and incubated with 2 mg of ICAM-1-coated beads at 37 °C on a rotating shaker for 60 min. Afterwards, the beads were washed with 0.3 mL of SELEX buffer prewarmed to 37 °C and bound aptamers were immediately eluted through denaturation at 94 °C for 3 min in nuclease free water. Half of the solution was used for emulsion PCR (ePCR), a technique designed to amplify and enrich individual DNA sequences within a reaction, prepared by mixing aqueous PCR components with the oil emulsion system supplied in the kit under controlled temperature. PCR reactions were prepared in a final volume of 50 μL containing 1× PCR buffer, 1.5 mM MgCl₂, 0.01 mg/mL acetylated bovine serum albumin (BSA; EURx, Poland, cat. n°: E4020), 400 μM of each dNTP, and 4 μM of each forward and reverse primer. Thermostable DNA polymerase (EURx Taq DNA polymerase, 5 U/µL; cat. n°: E2500) was added at a final amount of 1.25 U per reaction. Amplification was carried out using an Applied Biosystems 96-well thermal cycler (Fisher Scientific, USA, cat. n°: 12333653) under the following conditions: initial denaturation at 95 °C for 2 min, 20 cycles of 95 °C for 30 s, 55 °C for 60 s and 72 °C for 3 min, followed by a final extension at 72 °C for 5 min. After amplification, emulsions were broken and DNA was purified using spin columns. Final elution was performed by heating at 94 °C for 3 min. Amplification efficiency was verified by electrophoresis on a 3% agarose gel. DNA concentration was further quantified fluorometrically using the Quantifluor ONE ds DNA kit (Promega, USA, cat. n°: E4871) on a Quantus^TM^ Fluorometer (Promega, USA, cat. n°: E6150). A negative selection step was introduced after the first round to remove non-specifically binding sequences. Briefly, 50 pmol of enriched ssDNA from the first round were incubated with 0.2 mg of unloaded (empty) streptavidin-coated beads in SELEX buffer for 30 min at 37 °C on a rotating shaker. After incubation, bead-bound sequences were discarded, and the supernatant containing unbound DNA was recovered and subsequently incubated with 1 mg of ICAM-1-coated beads for the positive selection step. Positive selection cycles were repeated for a total of 10 rounds. Selection stringency was progressively increased by reducing incubation time (from 60 to 30 min), increasing wash volume (from 300 to 750 µL), and increasing the number of wash steps (from 1 to 3 washes per cycle), in order to enrich for high-affinity and high-specificity aptamers (Table S2). From the 4^th^ round onward, a fraction of the amplified DNA was set aside for Next-Generation Sequencing (NGS) using the Illumina MiSeq platform. Sequencing data were analysed using FastQC software for quality control and assessment of sequence enrichment (https://www.bioinformatics.babraham.ac.uk/projects/fastqc/).

### Single cycle SELEX assay

The binding ability of each aptamer identified to the ICAM-1 29-50 peptide was evaluated with a single cycle of the SELEX process. To this end, each synthesized aptamer (Merck Oligo, UK) was diluted in SELEX buffer at the final concentration of 0.2 µM, subjected to the denaturation-renaturation procedure in a final volume of 250 μL; 0.1 mg beads coated with the ICAM-1 peptide or, alternatively, the same amount of uncoated beads (empty), were added to the denatured-renatured aptamers. The mixture was incubated for 30 min at 37 °C with gentle agitation. After incubation, beads were washed twice with 1 mL SELEX buffer prewarmed to 37 °C to remove unbound sequences. The binder sequences were finally eluted from the beads through denaturation at 94 °C for 3 min in 100 μL of nuclease free water. To quantitatively evaluate the amount of bound material, 5 μL of the solution was assayed in a RT-PCR reaction in triplicate using SYBR green and GoTaq qPCR Master Mix (Promega, USA, cat. n°: A6001) according to the manufacturer’s protocol. The reaction was carried out in a 7500 Real-Time PCR System (Applied Biosystems, USA) with the following thermal profile: 2 min at 95 °C, then 40 cycles of 15 s at 95 °C, 1 min at 60 °C using the same primers used for the ePCR. Data were normalized based on Ct values, comparing for each aptamer the level of peptide-coated beads vs empty beads.

### Surface plasmon resonance (SPR) binding analysis

Binding of selected aptamers (I1-I8) to the ICAM-1 29-50 peptide was evaluated by surface plasmon resonance (SPR) using a Biacore 1K instrument (Cytiva) as previously described [39]. The biotinylated peptide was immobilized on a streptavidin-coated sensor chip according to the manufacturer’s instructions. Aptamers were injected at increasing concentrations (10, 30, and 100 nM) in PBS buffer at a flow rate of 10 μL/min at 25 °C. Binding responses were recorded, and equilibrium values were used to generate binding curves. Apparent dissociation constants (K_D_) were determined by fitting the data with a sigmoidal model.

### Aptamer structure prediction and molecular docking

To predict the most probable three-dimensional (3D) conformations of SELEX-enriched DNA sequences, the 3dDNA software (Xiao Lab) was used [40]. This tool employs libraries of DNA and RNA structural templates to model DNA 3D conformations based on secondary structure information. For each predicted secondary structure, up to five 3D models were generated by 3dDNA and ranked according to their free energy. The lowest-energy model was selected for subsequent docking analysis. Molecular docking was performed using the HADDOCK 2.4 web server [25]. The input structures consisted of the predicted aptamer models and the ICAM-1 chain extracted from the ICAM-1–HRV-A16 complex (PDB: 1D3E). Based on previous structural data [23], lysine at position 29 (K29) of ICAM-1 was defined as the active residue, while surrounding residues were automatically assigned as passive residues. Docking was carried out using the standard HADDOCK protocol, including rigid-body docking followed by semi-flexible refinement, generating an initial ensemble of thousands of poses that were subsequently refined and clustered.

### Cell culture

H1-HeLa cell line represents the standard for studying the biology of HRVs [41]. Cells were obtained from ATCC (LGC, Italy, cat. n°: CRL-1958) and maintained in Dulbecco’s Modified Eagle Medium (DMEM) High Glucose (Gibco™, Thermo Fisher Scientific, USA, cat. n°: 11965092) supplemented with 10% fetal bovine serum (FBS) and incubated at 37 °C in a humidified atmosphere containing 5% CO_2_. Cells were subcultured every 2-3 days.

### Viral strain

HRV-A16 (Human Rhinovirus A 16) strain 11757 was purchased from ATCC (LGC, Italy, cat. n°: VR-283). Virus was propagated in H1-HeLa cells in DMEM with 2% FBS at 33 °C and 5% CO_2_. Cells were infected at 33 °C because this temperature mimics the conditions of the human upper respiratory tract, the primary site of infection [42]. Following propagation, viral titres were determined using TCID_50_ assay and expressed as TCID_50_/ml. Aliquots of virus were prepared and stored at -80 °C to prevent repeated freeze-thaw cycles. All procedures involving viruses were handled in accordance with Biosafety level 2 (BSL-2) guidelines.

### Cell infection and antiviral assay of aptamers

H1-HeLa cells were seeded in 24-well plates at a density of 1 x 10^5^ cells per well one day prior to infection. On the day of infection, cells were washed with PBS, and medium was replaced with treatments or vehicle. Aptamers I4, I5 and I8 (Merck, UK) were prepared as previously described through a denaturation (94 °C 3 min) followed by renaturation on ice for 5 min to ensure proper folding. The process was performed in SELEX buffer 1X formulated without Tween-20 to ensure cell compatibility. Following renaturation, aptamers or vehicle were immediately added to the cells and incubated for 7 min at 37 °C. Pleconaril (Merck, USA, cat. n°: SML0307) was included as a positive control for antiviral activity [43]. Cells were pretreated with aptamers/pleconaril prior to viral exposure to allow binding to cell surface targets. Following pretreatment, cells were infected with HRV-A16 at a multiplicity of infection (MOI) of 0.1. Mock-infected controls were treated with medium only and processed in parallel. Cells were incubated at 33 °C for 1 hour on a rotating shaker to favor viral binding. Subsequently, cells were washed to remove unbound viruses and supplemented with fresh medium to support viral replication. Infection was allowed to proceed for 72 hours at 33 °C. At the end of the incubation period, culture supernatants were collected for viral RNA quantification and viral titer determination by TCID_50_ assay.

### TCID_50_ assay

Viral titers were determined by TCID_50_ assay. Serial ten-fold dilutions of viral stock or infection samples were added to confluent H1-HeLa cell monolayers in 96-well plates (six replicates per dilution) and incubated at 33 °C, 5% CO_2_ for 72 hours. Infection was assessed by virus-induced cytopathic effect (CPE) after crystal violet staining, and TCID₅₀/mL was calculated using the Reed and Muench method [44]. All procedures were performed under BSL-2 conditions.

### Viral RNA extraction

Viral RNA was extracted from media collected 72 hours post-infection (p.i.) using the QiAmp Viral RNA Mini Kit (Qiagen, Germany, cat. n°: 52904), according to the manufacturer’s instructions. Briefly, following clarification by centrifugation, samples were lysed under BSL-2 conditions using AVL buffer supplemented with carrier RNA, to ensure complete disruption of viral particles and enhance RNA recovery. Viral RNA selectively bound to the QiAmp silica membrane, followed by two washing steps (AW1 and AW2 buffers). Viral RNA was eluted in 50 µL of nuclease-free water and stored at -80 °C until use.

### Real-Time PCR

For viral RNA quantification, a commercial multiplex PCR assay (Fast Track Diagnostics respiratory pathogen 21, FTD) (Siemens Healthiness Company, Luxembourg), was used following the manufacturer’s protocol. Reactions were performed on a CFX96 Real-Time PCR Detection System (Bio-Rad, USA). The thermal profile consisted of an initial reverse transcription step at 50 °C for 15 min, followed by a denaturation step at 94 °C for 1 min. Amplification was carried out over 39 cycles of denaturation (94 °C for 1 min), annealing (94 °C for 8 s), and extension (60 °C for 1 min), with fluorescence signals collected during the annealing phase for real-time monitoring.

## Declarations

### Author contributions

CRediT: JD: Conceptualization, Investigation, Methodology, Project administration, Supervision, Validation, Visualization, Writing – original draft, Writing – review & editing; CC: Investigation, Validation, Writing – review & editing; SA: Investigation, Validation, Writing – review & editing; MR: Resources; VP: Investigation, Validation; TC: Investigation, Methodology, Validation, Visualization, Writing – review & editing; LC: Investigation, Validation; SR: Investigation, Validation; SDC: Investigation, Validation; AP: Investigation, Visualization, Writing – review & editing; CT: Resources, Supervision, Writing – review & editing; MB: Methodology, Resources, Supervision, Writing – review & editing; TB: Resources, Supervision, Writing – review & editing; AR: Methodology, Supervision, Writing – review & editing; AV: Methodology, Resources, Supervision; EB: Conceptualization, Methodology, Resources, Supervision, Writing – review & editing; EBo: Methodology, Supervision, Writing – review & editing; PC: Conceptualization, Funding acquisition, Methodology, Resources, Supervision, Validation, Writing – review & editing

### Corresponding author

Correspondence to

### Availability of data and materials

The data that support the fundings of this study are available from corresponding author upon reasonable request.

## Supplementary methods

### Supplementary structural and molecular dynamics analysis

a detailed structure-guided analysis of the HRV–ICAM-1 interaction was performed starting from the crystal structure (PDB: 1D3E). Missing hydrogen atoms were added using VEGA ZZ and the system was energy-minimized in two steps with NAMD3 (CHARMM36m force field, dielectric constant = 20): 30,000 steps with backbone restraints followed by 10,000 steps without constraints. The optimize complex was analysed by calculating non-bonded interaction energies (Lennard-Jones and distance-dependent Coulomb terms), which were decomposed at the residue level to identify ICAM-1 regions contributing to binding. This analysis, together with previous structural data, guided the selection of peptide candidates spanning the HRV-binding interface centered on Lys29. Among the candidates, peptide 4 (ICAM-1 residues 29–50) was selected based on a combination of favourable interaction energy, binding efficiency, and enrichment in positively charged residues, supporting its potential compatibility with nucleic acid ligands. To further assess its conformational behaviour, peptide 4 was subjected to molecular dynamics simulations in explicit solvent using NAMD3. The system was neutralized, solvated in a 45 Å cubic water box, and minimized (20,000 steps), followed by heating (0–300 K, 100 ps), equilibration (NPT, 5 ns), pre-production (NVT, 1 ns), and a 200 ns production run (NVT, 1 fs timestep). The time evolution of Cα-RMSD (Fig. S1) showed a fluctuating profile (average 12.1 Å), indicating conformational flexibility with recurrent sampling of native-like states. Residue-wise Cα-RMSF analysis (Table S1) revealed higher flexibility at the termini (including Lys29) and a more stable central region corresponding to the HRV-binding interface. The solvent-accessible polar surface area remained relatively stable throughout the simulation (∼4% variation), indicating preservation of key physicochemical features relevant for nucleic acid interaction. Overall, peptide 4 (ICAM-1 29-50) samples a dynamic conformational ensemble while retaining bioactive-like structural and physicochemical properties, supporting its use as a mimic of the ICAM-1 binding interface in SELEX experiments.

---

**Fig. S1.**
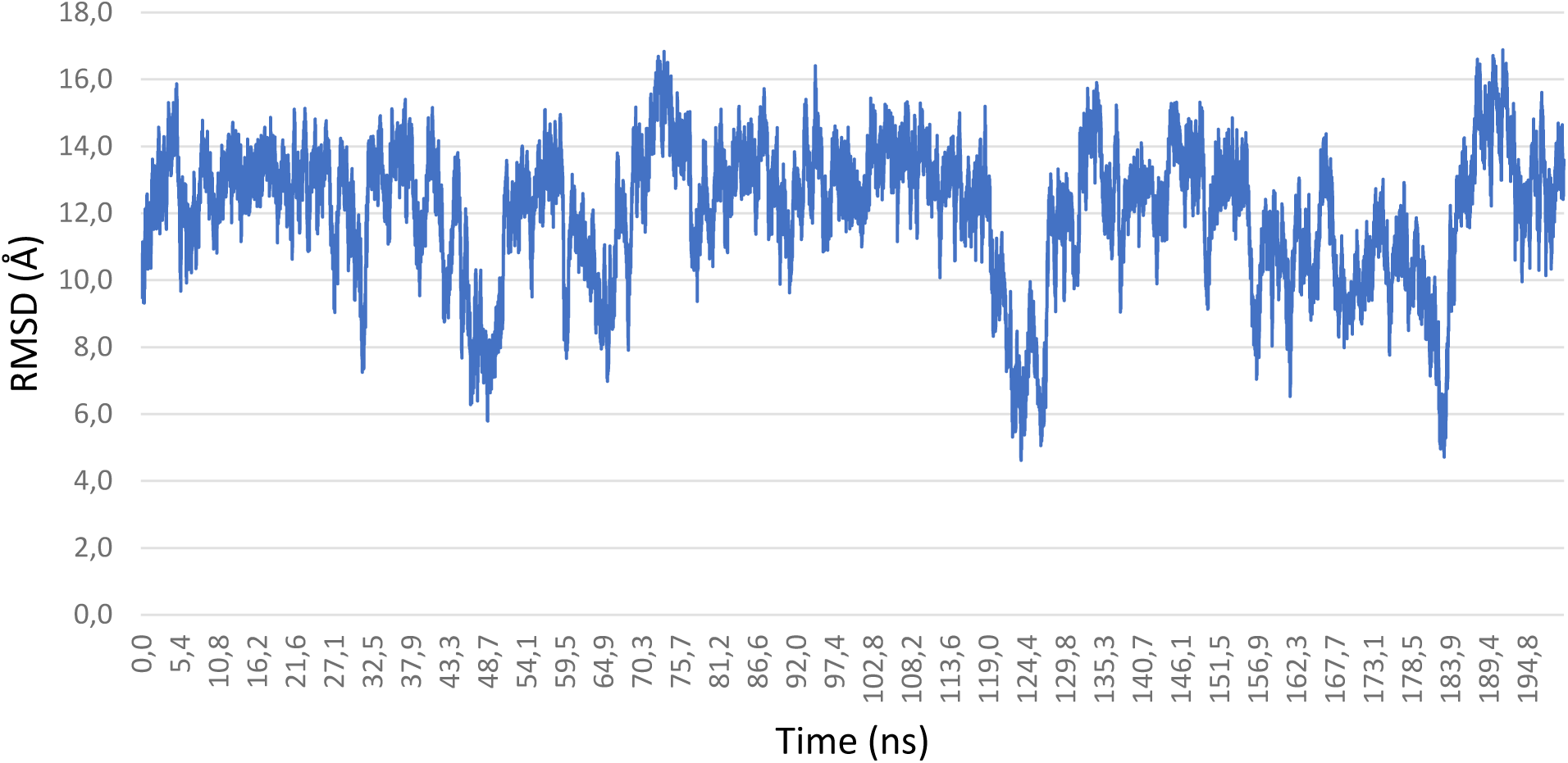
Time evolution of the Cα-RMSD of ICAM-1 29-50 peptide during the 200 ns molecular dynamics simulation. RMSD values (Å) were calculated with respect to the crystallographic conformation extracted from the HRV–ICAM-1 complex (PDB:1D3E) after structural alignment. The trajectory shows a broad distribution of RMSD values (4.6–16.9 Å; average 12.1 Å), indicative of significant conformational flexibility in solution. Recurrent decreases toward lower RMSD values are observed throughout the simulation, highlighting the ability of the peptide to repeatedly sample conformations resembling the native bound state.

**Fig. S2.**
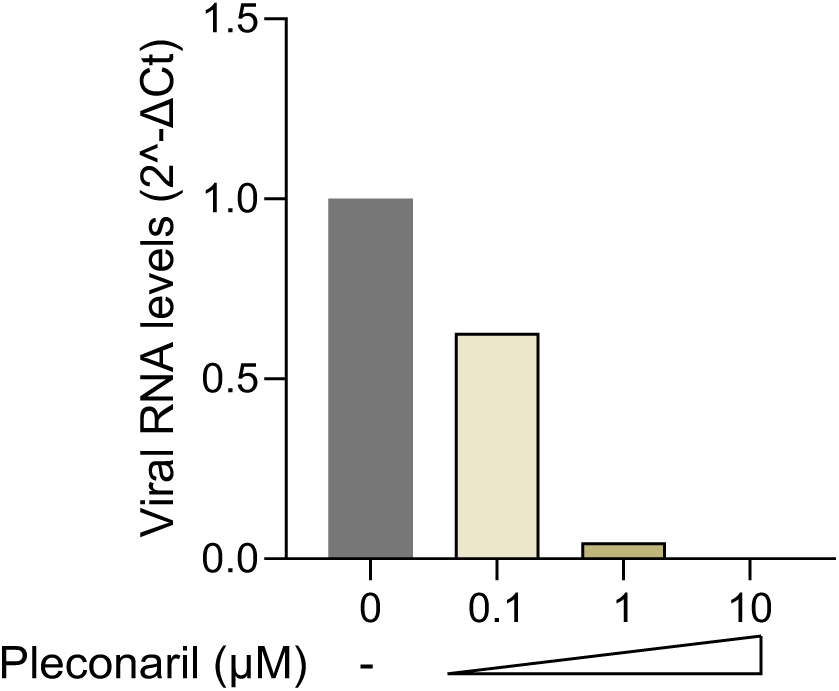
Dose-response effect of Pleconaril. The VP1-inhibitor Pleconaril was used as a positive control for infection inhibition. H1-HeLa cells were infected with HRV-A16 (MOI 0.1) and treated with increasing concentration of Pleconaril (0, 0.1, 1 and 10 µM). Viral RNA levels in culture supernatants were quantified 72 hours post infection by RT-PCR.

**Table S1.** Residue-wise Cα root mean square fluctuation (RMSF) of peptide 4 (residues 29–50) calculated over the 200 ns molecular dynamics simulation. RMSF values (Å) were obtained after alignment of the trajectory and reflect the flexibility of individual residues around their average positions. The analysis reveals a heterogeneous flexibility profile, with higher fluctuations at the N- and C-terminal regions (notably Lys29, Arg49, and Lys50, in red) and comparatively lower fluctuations in the central segment of the peptide (minimum at Glu41, in blue), indicating the presence of a relatively more stable core corresponding to the HRV-binding interface.

| Residue | Number | RMSF |
| --- | --- | --- |
| LYS | 29 | 7.89 |
| LEU | 30 | 5.88 |
| LEU | 31 | 4.58 |
| GLY | 32 | 4.18 |
| ILE | 33 | 4.17 |
| GLU | 34 | 3.42 |
| THR | 35 | 4.03 |
| PRO | 36 | 4.44 |
| LEU | 37 | 3.59 |
| PRO | 38 | 3.83 |
| LYS | 39 | 3.48 |
| LYS | 40 | 3.27 |
| GLU | 41 | 3.16 |
| LEU | 42 | 3.68 |
| LEU | 43 | 3.81 |
| LEU | 44 | 3.68 |
| PRO | 45 | 4.09 |
| GLY | 46 | 4.02 |
| ASN | 47 | 4.31 |
| ASN | 48 | 4.86 |
| ARG | 49 | 6.08 |
| LYS | 50 | 7.82 |

**Table S2.** SELEX protocol applied. The table summarizes the conditions applied for each SELEX round. The protocol progressively increases selection stringency by stepwise adjustment of binding and washing parameters: starting with relaxed conditions in Round 1 (longer incubation, minimal washing) to favor initial binding and gradually introducing shorter incubation times and multiple wash steps in later rounds to enrich higher-affinity binders.

| ICAM1 | Input | Reaction parameters |  |  |  |
| --- | --- | --- | --- | --- | --- |
| Round | Input DNA [μg] | Beads + target peptide [μg][pmol] | Binding incubation time [min] | Washing step (mL) | Number of PCR cycles |
| 1 | 100 | 2000\|800 | 60 | 0.3 | 20 |
| Negative selection |  |  |  |  |  |
| 2 | 0.6 | 1100\|440 | 60 | 1 x 0.38 | 20 |
| 3 | 0.45 | 1000\|400 | 60 | 1 x 0.45 | 20 |
| 4 | 0.50 | 1000\|400 | 50 | 2 x 0.30 | 20 |
| 5 | 0.80 | 1000\|400 | 50 | 2 x 0.38 | 20 |
| 6 | 0.45 | 1000\|400 | 50 | 3 x 0.40 | 20 |
| 7 | 0.45 | 1000\|400 | 40 | 3 x 0.50 | 20 |
| 8 | 0.39 | 1000\|400 | 40 | 3 x 0.60 | 20 |
| 9 | 0.40 | 1000\|400 | 30 | 3 x 0.70 | 20 |
| 10 | 0.3 | 800\|320 | 30 | 3 x 0.75 | 20 |

## References

[1] Jacobs SE, Lamson DM, Kirsten S, Walsh TJ. Human rhinoviruses. Clin Microbiol Rev 2013;26:135–62. 10.1128/CMR.00077-12.

[2] Rollinger JM, Schmidtke M. The human rhinovirus: Human-pathological impact, mechanisms of antirhinoviral agents, and strategies for their discovery. Med Res Rev 2011;31:42–92. 10.1002/med.20176.

[3] Liu J, Wang W, Cao K, Ren Z, Fu X, Chen Y, et al. Epidemiology and clinical characteristics of human rhinovirus in hospitalized children and adolescents with acute respiratory infections: a longitudinal study in Shenzhen, China (2019–2024). Virology Journal 2025;22. 10.1186/s12985-025-02901-9.

[4] Kieninger E, Fuchs O, Latzin P, Frey U, Regamey N. Rhinovirus infections in infancy and early childhood. European Respiratory Journal 2013;41:443–52. 10.1183/09031936.00203511.

[5] Salim S, Celiloglu H, Tayyab F, Malik ZA. Seasonal Prevalence of Respiratory Pathogens Among Children in the United Arab Emirates: A Multicenter Cross-Sectional Study in the Pre-COVID-19 Era. Cureus 2023. 10.7759/cureus.45204.

[6] Wedzicha JA, Singh R, Mackay AJ. Acute COPD Exacerbations. Clin Chest Med 2014;35:157–63. 10.1016/J.CCM.2013.11.001.

[7] Kennedy JL, Pham S, Borish L. Rhinovirus and Asthma Exacerbations. Immunol Allergy Clin North Am 2019;39:335–44. 10.1016/j.iac.2019.03.003.

[8] Bhutani M, Müllerová H, Patel D, Barjaktarevic I, Loke WJ, Pollack M, et al. Disease burden and health-related outcomes of patients discharged from hospital following a COPD exacerbation in the United States. Respir Med 2025;248:108337. 10.1016/J.RMED.2025.108337.

[9] Sánchez-Ramos J, Leonardo García-León M, Bautista-Carbajal P, Alfonso Salazar-Soto L, Noyola DE, Susana Juárez-Tobías M, et al. Academic Editors: Maria Pokorska-’ Spiewak and Magdalena Rutkowska Genotypic Diversity of Human Rhinovirus in Children with Pneumonia Before and During the COVID-19 Pandemic in Mexico 2025;14:1236. 10.3390/pathogens.

[10] Jackson DJ, Gern JE. Rhinovirus Infections and Their Roles in Asthma: Etiology and Exacerbations. Journal of Allergy and Clinical Immunology: In Practice 2022;10:673–81. 10.1016/j.jaip.2022.01.006.

[11] Halabi KC, Stockwell MS, Alba L, Vargas C, Reed C, Saiman L. Clinical and socioeconomic burden of rhinoviruses/enteroviruses in the community. Influenza Other Respir Viruses 2022;16:891–6. 10.1111/irv.12989.

[12] Palmenberg AC, Gern JE. Classification and evolution of human rhinoviruses. Methods in Molecular Biology 2015;1221. 10.1007/978-1-4939-1571-2_1.

[13] Casanova V, Sousa FH, Stevens C, Barlow PG. Antiviral therapeutic approaches for human rhinovirus infections. Future Virol 2018;13:505–18. 10.2217/fvl-2018-0016.

[14] Jacobs SE, Lamson DM, Kirsten S, Walsh TJ. Human rhinoviruses. Clin Microbiol Rev 2013;26:135–62. 10.1128/CMR.00077-12.

[15] Waman VP, Kolekar PS, Kale MM, Kulkarni-Kale U. Population structure and evolution of rhinoviruses. PLoS One 2014;9. 10.1371/journal.pone.0088981.

[16] MacArthur RD, Novak RM. Maraviroc: The first of a new class of antiretroviral agents. Clinical Infectious Diseases 2008;47:236–41. 10.1086/589289.

[17] Iacob SA, Iacob DG. Ibalizumab Targeting CD4 Receptors, An Emerging Molecule in HIV Therapy. Front Microbiol 2017;8. 10.3389/fmicb.2017.02323.

[18] Coultas JA, Cafferkey J, Mallia P, Johnston SL. Experimental antiviral therapeutic studies for human rhinovirus infections. J Exp Pharmacol 2021;13:645–59. 10.2147/JEP.S255211.

[19] Nimjee SM, White RR, Becker RC, Sullenger BA. Aptamers as Therapeutics. Annu Rev Pharmacol Toxicol 2017;57:61–79. 10.1146/ANNUREV-PHARMTOX-010716-104558.

[20] Villa A, Brunialti E, Dellavedova J, Meda C, Rebecchi M, Conti M, et al. DNA aptamers masking angiotensin converting enzyme 2 as an innovative way to treat SARS-CoV-2 pandemic. Pharmacol Res 2022;175:105982. 10.1016/J.PHRS.2021.105982.

[21] Keefe AD, Pai S, Ellington A. Aptamers as therapeutics. Nat Rev Drug Discov 2010;9:537–50. 10.1038/NRD3141.

[22] Kaur H, Bruno JG, Kumar A, Sharma TK. Aptamers in the Therapeutics and Diagnostics Pipelines. Theranostics 2018;8:4016–32. 10.7150/thno.25958.

[23] Kolatkar PR, Bella J, Olson NH, Bator CM, Baker TS, Rossmann MG, et al. Structural studies of two rhinovirus serotypes complexed with fragments of their cellular receptor. vol. 18. 1999.

[24] Bella J, Kolatkar PR, Marlor CW, Greve JM, Rossmann MG. The structure of the two amino-terminal domains of human ICAM-1 suggests how it functions as a rhinovirus receptor and as an LFA-1 integrin ligand. vol. 95. 1998.

[25] Honorato R V., Trellet ME, Jiménez-García B, Schaarschmidt JJ, Giulini M, Reys V, et al. The HADDOCK2.4 web server for integrative modeling of biomolecular complexes. Nat Protoc 2024;19:3219–41. 10.1038/s41596-024-01011-0.

[26] Nava G, Zanchetta G, Giavazzi F, Buscaglia M. Label-free optical biosensors in the pandemic era 2022;11:4159–81. doi:10.1515/nanoph-2022-0354.

[27] Ren Z, Shen C, Peng J. Status and Developing Strategies for Neutralizing Monoclonal Antibody Therapy in the Omicron Era of COVID-19. Viruses 2023;15. 10.3390/v15061297.

[28] Low ZY, Farouk IA, Lal SK. Drug repositioning: New approaches and future prospects for life-debilitating diseases and the COVID-19 pandemic outbreak. Viruses 2020;12. 10.3390/v12091058.

[29] Muthukutty P, MacDonald J, Yoo SY. Combating Emerging Respiratory Viruses: Lessons and Future Antiviral Strategies. Vaccines (Basel) 2024;12. 10.3390/vaccines12111220.

[30] Esneau C, Duff AC, Bartlett NW. Understanding Rhinovirus Circulation and Impact on Illness. Viruses 2022;14. 10.3390/v14010141.

[31] Shukla SD, Shastri MD, Vanka SK, Jha NK, Dureja H, Gupta G, et al. Targeting intercellular adhesion molecule-1 (ICAM-1) to reduce rhinovirus-induced acute exacerbations in chronic respiratory diseases. Inflammopharmacology 2022;30:725–35. 10.1007/s10787-022-00968-2.

[32] Traub S, Nikonova A, Carruthers A, Dunmore R, Vousden KA, Gogsadze L, et al. An Anti-Human ICAM-1 Antibody Inhibits Rhinovirus-Induced Exacerbations of Lung Inflammation. PLoS Pathog 2013;9. 10.1371/journal.ppat.1003520.

[33] Zhou J, Rossi J. Aptamers as targeted therapeutics: Current potential and challenges. Nat Rev Drug Discov 2017;16:181–202. 10.1038/nrd.2016.199.

[34] Gao S, Zheng X, Jiao B, Wang L. Post-SELEX optimization of aptamers. Anal Bioanal Chem 2016;408:4567–73. 10.1007/s00216-016-9556-2.

[35] Yu H, Zhu J, Shen G, Deng Y, Geng X, Wang L. Improving aptamer performance: key factors and strategies. Microchimica Acta 2023;190:255. 10.1007/s00604-023-05836-6.

[36] Pedretti A, Mazzolari A, Gervasoni S, Fumagalli L, Vistoli G. The VEGA suite of programs: An versatile platform for cheminformatics and drug design projects. Bioinformatics 2021;37:1174–5. 10.1093/bioinformatics/btaa774.

[37] Phillips JC, Hardy DJ, Maia JDC, Stone JE, Ribeiro J V., Bernardi RC, et al. Scalable molecular dynamics on CPU and GPU architectures with NAMD. Journal of Chemical Physics 2020;153. 10.1063/5.0014475.

[38] Brooks BR, Brooks CL, Mackerell AD, Nilsson L, Petrella RJ, Roux B, et al. CHARMM: The biomolecular simulation program. J Comput Chem 2009;30:1545–614. 10.1002/jcc.21287.

[39] Muniz MI, Carzaniga T, Nava G, Casiraghi L, Giana D, Rocca S, et al. Sequence optimization of a DNA aptamer inhibiting COVID-19 infection guided by analysis of secondary structure distribution. Comput Struct Biotechnol J 2026;31:130–42. 10.1016/j.csbj.2025.12.010.

[40] Zhang Y, Xiong Y, Xiao Y. 3dDNA: A Computational Method of Building DNA 3D Structures. Molecules 2022;27. 10.3390/molecules27185936.

[41] Lee W-M, Chen Y, Wang W, Mosser A. Growth of Human Rhinovirus in H1-HeLa Cell Suspension Culture and Purification of Virions. In: Jans DA, Ghildyal R, editors. Rhinoviruses: Methods and Protocols, New York, NY: Springer New York; 2015, p. 49–61. 10.1007/978-1-4939-1571-2_5.

[42] Foxman EF, Storer JA, Fitzgerald ME, Wasik BR, Hou L, Zhao H, et al. Temperature-dependent innate defense against the common cold virus limits viral replication at warm temperature in mouse airway cells. Proc Natl Acad Sci U S A 2015;112:827–32. 10.1073/pnas.1411030112.

[43] Pevear DC, Tull TM, Seipel ME, Groarke JM. Activity of pleconaril against enteroviruses. Antimicrob Agents Chemother 1999;43. 10.1128/aac.43.9.2109.

[44] Reed LJ, Muench H. A Simple Method of Estimating Fifty per cent Endpoints. Am J Epidemiol 1938;27:493–7. 10.1093/oxfordjournals.aje.a118408.

